# TAK1 integrates the NLRP1 inflammasome into the innate immune response to double-stranded RNA

**DOI:** 10.64898/2026.01.23.701331

**Authors:** Miles R. Corley, Ayumu Hyodo, Hunter C. Toyoda, Marisa A. Yonemitsu, Amandine Chantharath, Jennifer L. Hyde, Patrick S. Mitchell

## Abstract

Innate immune recognition of double-stranded RNA (dsRNA) by germline-encoded receptors initiates antiviral defenses, including type I interferon (IFN) production. The inflammasome-forming sensor NLRP1 binds and is activated by dsRNA in a mitogen-activated protein kinase (MAPK) p38-dependent manner. How dsRNA initiates these events to induce NLRP1 inflammasome activation is unclear. Here we demonstrate that both exogenous and cellular dsRNA triggers NLRP1 inflammasome activation downstream of RIG-I/MDA5–MAVS and/or TLR3–TRIF signaling but is independent of type I IFN. In immortalized and primary human keratinocytes, we find that NLRP1 inflammasome activation by dsRNA, including during viral infection, requires the MAPK kinase kinase TAK1. Mechanistically, TAK1-dependent phosphorylation of the NLRP1 N-terminal disordered region is necessary and sufficient for inflammasome activation. Collectively, we reveal TAK1 as a novel activator of the NLRP1 inflammasome, functioning as a critical signaling hub linking NLRP1 to inflammatory responses in the context of viral infection and autoimmunity.

## INTRODUCTION

Virtually all organisms are subject to pathogen attack, driving the evolution of host strategies for the recognition of and response to infectious threats. In vertebrates, ‘self’ is distinguished from microbial ‘non-self’ through the innate immune recognition of microbe-specific structures called pathogen-associated molecular patterns (PAMPs) by germline-encoded pattern recognition receptors (PRRs), coordinating pathogen recognition with the initiation of host defenses^1,2^.

During viral infection, nucleic acids such as double-stranded RNA (dsRNA) are a primary target of innate immune recognition^3,4^. Vertebrates encode several dedicated dsRNA PRRs that initiate a wide range of host responses^5-8^. For example, retinoic acid-inducible gene I (RIG-I) and melanoma differentiation-associated protein 5 (MDA5) detect distinct features of cytosolic dsRNA, leading to the activation of mitochondrial antiviral signaling protein (MAVS), the production of type I interferon (IFN), and establishment of the antiviral state. Similarly, toll-like receptor 3 (TLR3) detects endosomal dsRNA and initiates IFN production via the TIR domain containing adaptor molecule 1 (TRIF). The recognition of dsRNA also triggers oligoadenylate synthetase (OAS)/ribonuclease L (RNase L)-induced mRNA degradation, protein kinase R (PKR)-induced host translation arrest, Z-DNA binding protein 1 (ZBP1)-induced necroptotic cell death, and other innate immune responses that play a critical role in host defense. Aberrant activation of dsRNA PRR pathways also contributes to human autoinflammatory diseases^9^. Thus, understanding the molecular mechanisms by which dsRNA triggers host responses is important for the development of therapeutic strategies that modulate inflammation in the context of infectious and autoinflammatory disease.

Inflammasomes are an important component of vertebrate innate immunity, with well-established roles in host defense against diverse pathogens^10-12^. Upon the detection of pathogens or noxious stimuli, inflammasome-forming sensors assemble into cytosolic immune complexes for the recruitment and activation of pro-inflammatory caspases (e.g. caspase-1 (CASP1)), which can occur either directly or through the adaptor ASC (apoptosis-associated speck-like protein containing a caspase activation and recruitment domain (CARD)). Canonical inflammasome assembly initiates inflammatory signaling via CASP1-dependent processing of interleukin (IL)-1β and IL-18. Activated CASP1 also cleaves the pore-forming protein Gasdermin D (GSDMD), which initiates pyroptotic cell death^13^. Although predominantly studied in the context of myeloid cells, inflammasomes are also expressed in epithelial and endothelial cells and play an important role in barrier defense^14,15^.

Many inflammasome-forming sensors function as classical PRRs wherein ligand binding induces a conformational change that drives inflammasome assembly and activation^10,11,16^. In contrast, the NLRP1 inflammasome can be activated by pathogen-associated activities^17,18^. NLRP1 consists of an N-terminal pyrin domain (PYD), a disordered region (DR), nucleotide binding domain (NBD), and leucine rich repeats (LRRs), which are separated from a C-terminal CARD by a function-to-find domain (FIIND). Autolytic processing of the FIIND^19-21^, which is comprised of ZU5 and UPA subdomains, results in the non-covalent association of the PYD-DR-NBD-LRR-ZU5 and UPA-CARD. This unique domain architecture permits NLRP1 to directly sense and be activated by the enzymatic function of pathogen-encoded virulence factors (i.e., effectors) that destabilize its N-terminus – an example of effector-triggered immunity^22-24^. For instance, the NLRP1-DR harbors substrate mimics of viral and bacterial proteases^25-32^. Proteolytic cleavage of the DR generates a neo-N-terminus that is marked for proteasomal degradation by the ‘N-end rule’ pathway, leading to the release and assembly of the C-terminal UPA-CARD and subsequent inflammasome activation^27,33,34^.

The NLRP1 inflammasome also indirectly senses pathogen-associated activities that disrupt host translation via its integration into the ribotoxic stress response (RSR)^35-38^. For example, *Diphtheria* toxin and *Pseudomonas aeruginosa* exotoxin A inhibit the eukaryotic elongation factor 2 (eEF2), resulting in ribosome stalling and collisions that induce ribotoxic stress. Similarly, Anisomycin (ANS), an antibiotic from *Streptomyces griseolus*, also induces the RSR. Each of these perturbations trigger NLRP1 inflammasome activation in a manner dependent on the ribosome-bound, mitogen-activated protein kinase (MAPK) kinase kinase MAP3K20 (also known as ZAKα). ZAKα is a sensor of ribosome collisions, and its activation initiates a signal transduction cascade that activates MAP2Ks and MAPKs, including p38^39,40^. In response to ribotoxic stress, ZAKα and p38 phosphorylation of the NLRP1-DR drives inflammasome activation^36,38^, implicating NLRP1 in the etiology of inflammatory skin pathologies from bacterial toxins or environmental insults like UVB irradiation^41-43^.

Recently, dsRNA was also found to induce NLRP1 inflammasome activation^44^. For example, arthritogenic alphavirus infection or the synthetic dsRNA mimic polyinosinic:polycytidylic acid (poly(I:C)) triggers NLRP1 inflammasome activation and the maturation and release of IL-1β and IL-18 from human keratinocytes^36,44,45^. Intron-containing HIV-1 RNA also triggers NLRP1 inflammasome actication^46^. Mechanistically, NLRP1’s response to dsRNA requires two molecular events: 1) binding to dsRNA via a positively charged groove in its LRRs, akin to canonical PRRs^44^, and 2) activation of p38, reminiscent of RSR-triggered NLRP1 inflammasome activation^36,45^. However, the pathway leading to p38 kinase activation in response to dsRNA is unknown.

Here we demonstrate that RIG-I, MDA5, and/or TLR3 signaling is specifically required for the NLRP1 inflammasome response to dsRNA, but not other activating stimuli, to activate the MAP3K TAK1. We find that genetic deletion or chemical inhibition of TAK1 prevents NLRP1 inflammasome activation in human keratinocytes in response to poly(I:C) and alphavirus infection. Mechanistically, our findings are consistent with TAK1 activating NLRP1 in a manner dependent on MAP3K phosphomotifs in the NLRP1-DR. Moreover, TAK1 is sufficient to induce NLRP1 inflammasome activation in the absence of exogenous dsRNA. Together our findings suggest that MAP3Ks function as hubs that allow NLRP1 to indirectly sense pathogen perturbations to cellular homeostasis, including TAK1, which integrates the NLRP1 inflammasome into dsRNA innate immune responses. Our model implicates the NLRP1 inflammasome in both human antiviral immunity and dsRNA-associated inflammatory diseases.

## RESULTS

### Pattern recognition of dsRNA is required upstream of NLRP1 inflammasome activation

To evaluate the requirements for dsRNA-triggered NLRP1 inflammasome activation, we first attempted to reconstitute the NLRP1 inflammasome in 293T cells, which lack endogenous expression of inflammasome components. As we and others have shown previously, co-transfection of plasmids encoding human NLRP1, ASC, CASP1, and pro-IL-1β recapitulates NLRP1-dependent CASP1 processing of IL-1β in response to well-established NLRP1 activating stimuli^25,31-33^. For example, Val-boroPro (VbP), an inhibitor of the dipeptidyl peptidases DPP8 and DPP9, which form a ternary inhibitory complex with NLRP1 to prevent spurious release of the UPA-CARD^47,48^, robustly induces CASP1-dependent processing of pro-IL-1β to p17 in a NLRP1-dependent manner, as measured by an IL-1 bioassay or immunoblotting (Fig. 1A and 1B). Similarly, co-transfection with plasmids encoding the coxsackievirus B3 (CVB3) 3C protease or ZAKα (a proxy for ribotoxic stress) also results in NLRP1-dependent inflammasome activation, (Fig. 1A and 1B). In contrast, we found that transfected poly(I:C) failed to induce NLRP1 inflammasome activation (Fig. 1A and 1B), suggesting that NLRP1 binding to dsRNA is not sufficient for inflammasome activation.

**Fig. 1.**
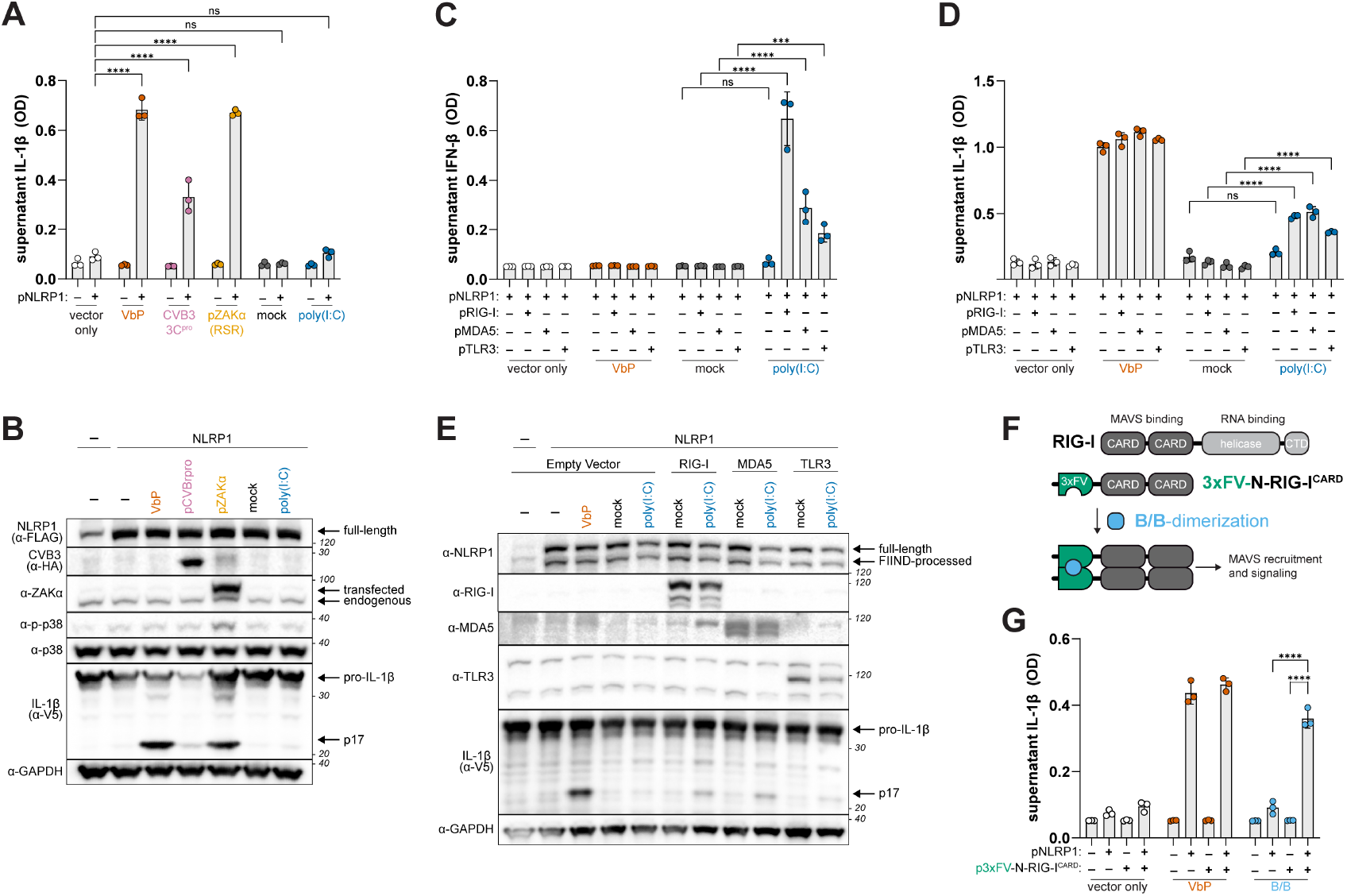
RNA sensors are required for NLRP1 activation by dsRNA. (**A-B**) Reconstitution of the NLRP1 inflammasome by co-transfecting 293T cells with plasmids encoding NLRP1, ASC, CASP1, and pro-IL-1β. Cells were treated with VbP (10μM), mock transfected, or co-transfected with HMW poly(I:C) (1μg/mL) or plasmids encoding ZAKα or CVB3 3C protease. Levels of bioactive IL-1β, reported as optical density (OD), were quantified using a HEK-Blue IL-1β reporter assay (A, see Methods). Immunoblotting of indicated proteins from lysates harvested 40-42 hours post-transfection (B). (**C-E**) NLRP1 inflammasome reconstituted 293T cells were co-transfected with empty vector, pRIG-I, pMDA5, or pTLR3 and treated with VbP (10μM), mock transfected, or transfected with HMW poly(I:C) (1μg/mL). Quantification of supernatant IL-1β levels (D) and immunoblotting (E) were carried out as previously described. Bioactive IFN-β levels were quantified using a HEK-Blue IFN reporter assay (C). (**F**) Schematic depicting drug (B/B) inducible murine RIG-I CARDs fused to FKBP12^F36V^ dimerizing domains (3xFV-N-RIG-I^CARD^). (**G**) NLRP1 inflammasome reconstituted 293T cells were co-transfected with either an empty vector or the 3xFV-N-RIG-I^CARD^ and treated with VbP (10μM) or B/B (10nM). IL-1β was quantified as in (A). Results are representative of two (A-B, E,), three (G), or four (C-D) independent experiments. Data are presented as mean ± SD. p-values were determined by two-way ANOVA with Tukey’s test. *p<0.05, **p<0.01, ***p<0.001, ****p<0.0001.

To account for this, we reasoned that 293T cells may lack a component upstream of p38 that is required for dsRNA-triggered NLRP1 inflammasome activation. Indeed, whereas ZAKα over-expression induced p38 phosphorylation and NLRP1 inflammasome activation, we were unable to detect phosphorylated p38 in cells transfected with poly(I:C) (Fig. 1B). We also noticed that poly(I:C) transfection of 293T cells did not lead to the production of type I IFN (Fig. 1C and fig. S1), which is likely due to low or undetectable levels of the dsRNA PRRs RIG-I, MDA5, or TLR3 (Fig. 1E, fig. S1). Indeed, we found that co-transfection of 293T cells with plasmids encoding RIG-I, MDA5, or TLR3 restored the production of type I IFN in response to poly(I:C) (Fig. 1C), also confirming the effective delivery of dsRNA to the host cytoplasm. Whereas the addition of dsRNA PRRs had no effect on VbP-induced NLRP1 inflammasome activation, surprisingly, the presence of RIG-I, MDA5, or TLR3 sensitized 293T cells to NLRP1-dependent inflammasome activation in response to poly(I:C) (Fig. 1D and 1E).

We next sought to decouple dsRNA recognition from RIG-I-like receptor (RLR) signaling in the context of NLRP1 inflammasome activation. To do so, we took advantage of a construct encoding tandem N-terminal RIG-I CARDs fused to FKBP12^F36V^ dimerization domains (3xFV-N-RIG-I^CARD^), which induces MAVS signaling and the production of type I IFN in the presence of the FKBP12-dimerizing small molecule AP20187 (B/B) (Fig. 1F)^49^. We found that co-transfection of 3xFV-N-RIG-I^CARD^ and inflammasome components in 293T cells induced inflammasome activation in a NLRP1-dependent manner in the presence but not absence of B/B (Fig. 1G), indicating that RIG-I–MAVS signaling is sufficient to activate the NLRP1 inflammasome in the absence of exogenous dsRNA.

To further examine the role of innate immune signaling in dsRNA-triggered NLRP1 inflammasome activation, we next generated genetic knockouts (KOs) of *NLRP1, TLR3*, and *MAVS* in the tert-immortalized human keratinocyte cell line N/TERT-2G (Fig. 2A-C), which we confirmed by western blotting (Fig. 2C and fig. S2A). As expected, the production of IFN-β by poly(I:C) was marginally affected in *TLR3* KO and *MAVS* KO cells but was ablated in *MAVS/TLR3* double KO (DKO) cells compared to WT N/TERT-2Gs, whereas supernatant IFN-β levels were unaffected in *NLRP1* KOs (Fig 2A, fig S2C). Similarly, supernatant IL-1β levels induced by poly(I:C) were similar across WT and *TLR3* KO and *MAVS* KO N/TERT-2Gs, consistent with prior reports^44,46^. However, poly(I:C)-triggered IL-1β release was abrogated in *MAVS*/*TLR3* DKO cells to the same extent as *NLRP1* KO cells (Fig. 2B, fig. S2B). Neither pro-IL-1β levels or inflammasome activation by VbP were affected in *MAVS* or *TLR3* single or double KO N/TERT-2Gs (Fig. 2B and 2C, fig. S2A), suggesting that MAVS and TLR3 signaling do not affect transcriptional priming and that their role in NLRP1 inflammasome activation is specific to dsRNA. Consistent with MAVS and TLR3 signaling being required to induce MAPK activation, p38 phosphorylation was inhibited in *MAVS*/*TLR3* DKO cells (Fig. 2C). Moreover, in addition to type I IFN production, MAVS or TRIF over-expression in 293T cells also induced NLRP1 inflammasome activation (Fig. 2D-F). Taken together, our findings indicate that dsRNA recognition and signaling by RIG-I, MDA5, or TLR3 is required to induce p38 phosphorylation upstream of NLRP1 inflammasome activation.

**Fig. 2.**
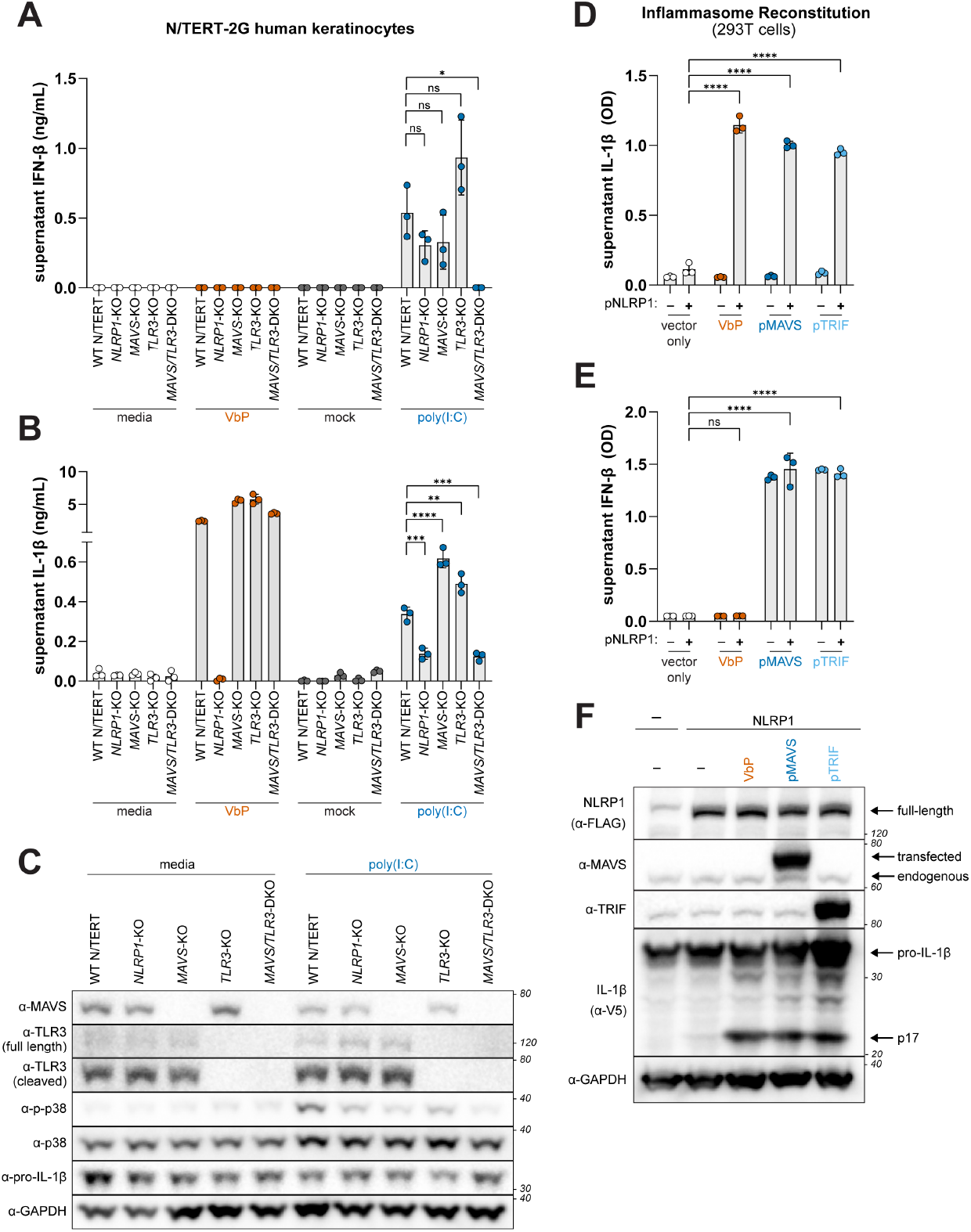
dsRNA recognition and innate immune signaling is required upstream of NLRP1 inflammasome activation. (**A-B**) WT and indicated single or double knockout (KO) N/TERT-2G cells were primed with 10ng/mL TNFα for 24 hours and then treated with VbP (10μM), mock transfected, or transfected with HMW poly(I:C) (1μg/mL). Supernatants were harvested 24 hours following the addition of activating stimuli and IL-1β (A) and IFN-β (B) were measured using an antibody-based immunoassay (see Methods). (**C**) Immunoblotting for indicated proteins from lysates harvested 6 hours following mock or poly(I:C) transfection of unprimed WT and indicated KO N/TERT-2G cells. (**D-F**) NLRP1 reconstituted 293T cells were either treated with VbP (10μM) or co-transfected with an empty vector, pMAVS, or pTRIF. IL-1β (D) and IFN-β (E) levels were quantified as described in Fig. 1. Immunoblotting for indicated proteins was carried out as described in (C) from lysates harvested 40-42 hours post-transfection (F). Results are representative of two (A-C) or three (D-F) independent experiments. Data are presented as mean ± SD. p-values were determined by one-way ANOVA with Dunnett’s test (A-B) or by two-way ANOVA with Tukey’s test (D-E). *p<0.05, **p<0.01, ***p<0.001, ****p<0.0001.

### Type I IFN signaling is not required for dsRNA-triggered NLRP1 inflammasome activation

We next sought to characterize the pathway by which RIG-I/MDA5–MAVS and TLR3– TRIF signaling induces NLRP1 inflammasome activation. We first considered if type I IFN is required for NLRP1 inflammasome activation. We found that pre-treatment of N/TERT-2G cells with the Janus kinase 1/2 inhibitor Ruxolitinib or an anti-IFN blocking antibody robustly inhibited the production of IFN-β and induction of the interferon-stimulated genes *MX1, IFIT1*, and *ISG15* by poly(I:C) (Fig. 3A and 3B), but had no effect on NLRP1-dependent IL-1β release from N/TERT-2G cells treated with VbP or poly(I:C) (Fig. 3C). Consistent with these observations, the level of phosphorylated p38 and supernatant IL-1β in response to VbP and poly(I:C) were comparable between WT and interferon regulatory factor 3 (*IRF3*) KO N/TERT-2G cells (Fig. 3D, 3E, and 3F). Furthermore, treatment with recombinant IFN-β (rIFN-β) did not induce nor augment inflammasome activation (Fig. 3C and 3F). We note that IFN-β did induce p38 phosphorylation (Fig. 3F), consistent with prior work indicating that p38 phosphorylation alone is not sufficient to activate NLRP1^36,37^. Taken together, our results indicate that the NLRP1 inflammasome response to dsRNA does not require type I IFN signaling.

**Fig. 3.**
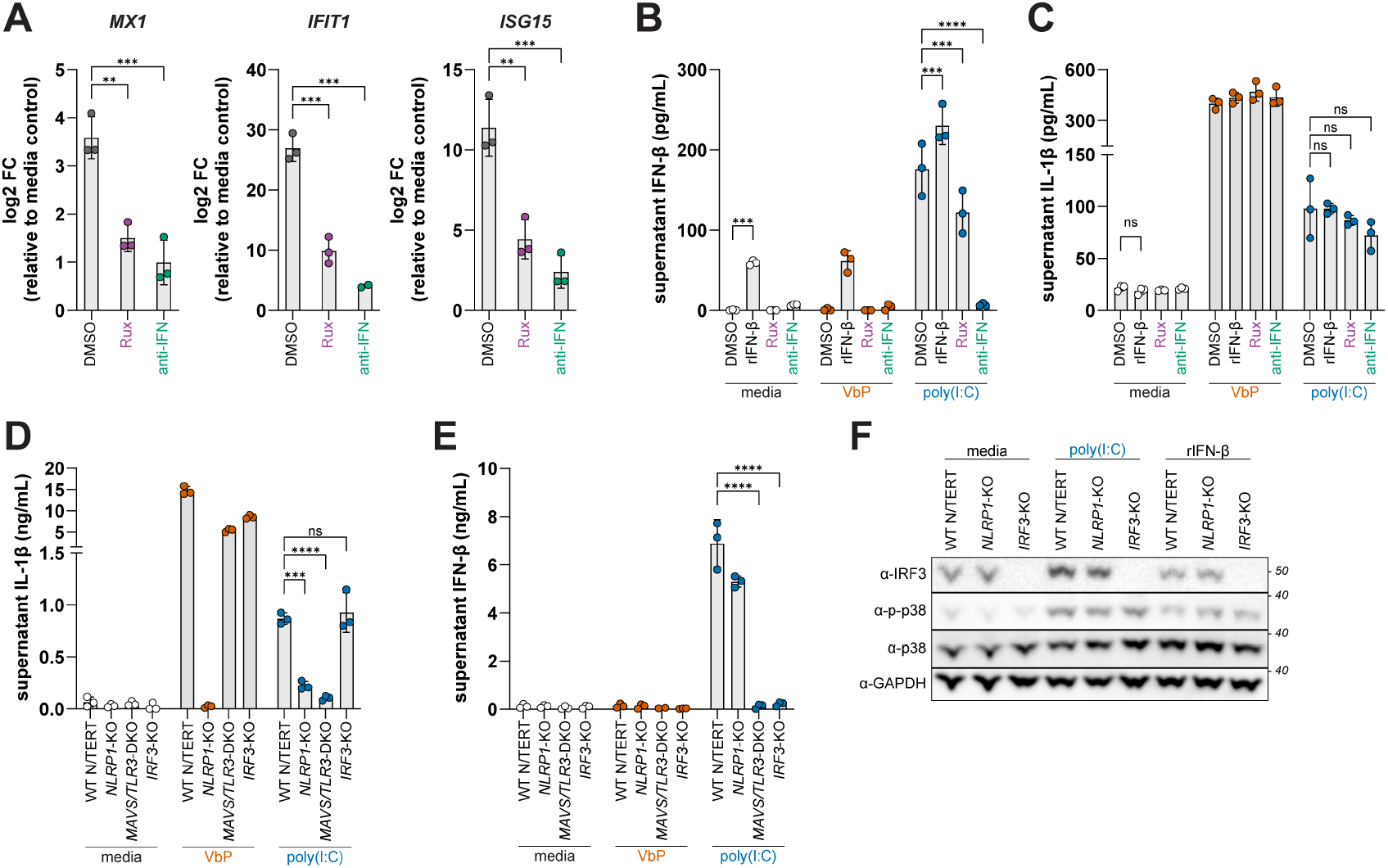
NLRP1 activation by dsRNA is independent of the type I IFN response. (**A-C**) N/TERT-2G cells were primed with 10ng/mL TNFα for 24 hours. Cells were incubated with either Ruxolitnib (Rux) (1μM) or IFN blocking antibody (anti-IFN) (1:50) for 1 hour prior to treatment with either VbP (10μM), recombinant IFN-β (rIFN-β), or HMW poly(I:C) (1μg/mL) and supernatants and cell pellets were harvested 10 hours post treatment. Expression of *MX1, IFIT1*, and *ISG15* in poly(I:C)-treated N/TERT-2G cells was determined by quantitative RT-PCR (A). IFN-β (B) and IL-1β (C) were quantified by ELISA. (**D-E**) WT and indicated KO N/TERT-2G cells were primed with 10ng/mL TNFα for 24 hours and treated with VbP (10μM) or HMW poly(I:C) (1μg/mL) for 24 hours. Supernatant IL-1β (D) and IFN-β (E) were quantified using an antibody-based immunoassay. (**F**) Immunoblotting from lysates generated from unprimed WT and indicated KO N/TERT-2G cells treated with VbP (10μM) or HMW poly(I:C) (1μg/mL) for 6 hours. Results are representative of two (A-F) independent experiments. Data are presented as mean ± SD. p-values were determined by one-way ANOVA with Dunnett’s test (A, D, and E) or two-way ANOVA with Sidak’s test (B-C). *p<0.05, **p<0.01, ***p<0.001, ****p<0.0001.

### TAK1 is required for dsRNA-triggered NLRP1 inflammasome activation

Our findings suggest that dsRNA induces p38 phosphorylation downstream of RIG-I/MDA5–MAVS and TLR3–TRIF signaling. To determine if ZAKα is a general requirement for phosphorylation-dependent modes of NLRP1 inflammasome activation, we treated WT and *MAP3K20* (ZAKα) KO N/TERT-2G cells with VbP, ANS, and poly(I:C). As expected, WT but not *MAP3K20* KOs were responsive to ANS-induced NLRP1 inflammasome activation. In contrast, *MAP3K20* KO cells responded normally to VbP and only modestly reduced poly(I:C)-triggered NLRP1 inflammasome activation (Fig. 4A). Interestingly, levels of phosphorylated p38 were also modestly reduced in poly(I:C) treated *MAP3K20* KO cells (Fig. 4B). Nevertheless, whereas ANS led to ZAKα phosphorylation (indicated by a mobility shift of ZAKα on a phostag gel), poly(I:C) did not, indicating that ZAKα does not respond to dsRNA and that RSR- and dsRNA-triggered NLRP1 inflammasome activation are distinct pathways upstream of p38.

**Fig. 4.**
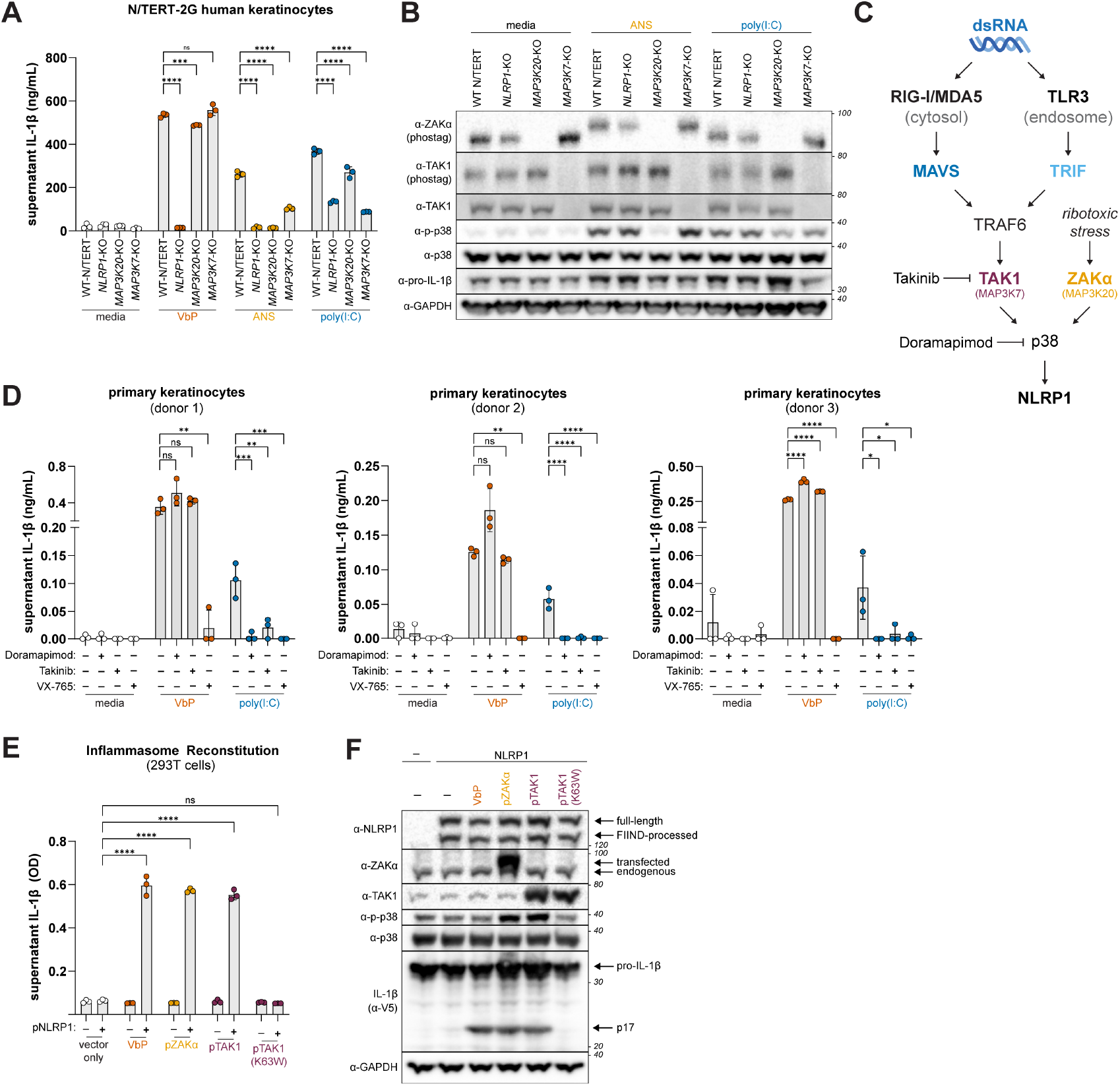
TAK1 is required for NLRP1 activation by dsRNA. (**A**) IL-1β levels were quantified by ELISA from supernatants of WT and indicated KO N/TERT-2G cells treated with VbP (10μM), ANS (1μM), or HMW poly(I:C) (1μg/mL) for 24 hours. (**B**) Immunoblotting for indicated proteins from lysates generated from N/TERT-2G cells treated as described in (A) 6 hours post-treatment. (**C**) Schematic depicting dsRNA activation of TAK1 and ribotoxic stress activation of ZAKa upstream of the NLRP1 inflammasome. (**D**) Primary keratinocytes from three independent donors were treated with Doramapimod (p38 inhibitor; 10μM) or Takinib (TAK1 inhibitor; 10μM) for 2 hours prior to treatment with VbP (10μM), ANS (1μM), or HMW poly(I:C) (1μg/mL) for 24 hours. Supernatant IL-1β levels were quantified by an antibody-based immunoassay. (**E-F**) NLRP1 reconstituted 293T cells were either treated with VbP (10μM) or co-transfected with pZAKα, pTAK1 WT, or pTAK1 K63W. Quantification of supernatant IL-1β using HEK-Blue IL-1β reporter cells (D). Immunoblotting for indicated proteins from lysates harvested 40-42 hours post-transfection (E). Results are representative of two (A-B) or three (D-F) independent experiments. Data are presented as mean ± SD. p-values were determined by two-way ANOVA with Sidak’s test (A, E) or by one-way ANOVA with Dunnett’s test (D) using GraphPad PRISM 10. *p<0.05, **p<0.01, ***p<0.001, ****p<0.0001.

We next considered a role for the MAP3K TAK1, which is activated downstream of both RIG-I/MDA5–MAVS and TLR3–TRIF pathways^50,51^. TAK1 also functions as a critical junction downstream of both the TNFα and IL-1 receptors to promote NF-кB signaling^52,53^, which could modulate inflammasome responses to dsRNA in other ways. However, we found that WT and *MAP3K7* (TAK1) KO cells respond similarly to VbP (Fig. 4A and fig. S3A), and we did not observe differences in the level of pro-IL-1β between WT and *MAP3K7* KO keratinocytes (Fig. 4B and fig. S3B). Pathogen inhibition of TAK1 can also induce pyroptosis of myeloid cells via CASP8 cleavage of GSDMD^54,55^. However, in both N/TERT-2G and primary human keratinocytes, neither *MAP3K7* KO nor treatment with the TAK1 inhibitor Takinib resulted in an increase of supernatant IL-1β (Fig. 4A and 4D). We thus hypothesized that TAK1 plays an analogous role to ZAKα in dsRNA-triggered NLRP1 inflammasome activation (Fig. 4C). Consistent with that possibility, we found that *MAP3K7* KO N/TERT-2G cells were unresponsive to poly(I:C) but responded to VbP comparably to WT cells (Fig. 4A; fig. S3A). Interestingly, ANS-induced NLRP1 inflammasome activation was modestly reduced in *MAP3K7* KO NTERT-2G cells (Fig. 4A), suggesting crosstalk between MAP3Ks in phosphorylation-dependent modes of NLRP1 inflammasome activation^45^.

To further assess the role of TAK1 in dsRNA-triggered NLRP1 inflammasome activation, we treated primary human keratinocytes from three independent donors with poly(I:C) in the presence or absence of Takinib, the p38 inhibitor Doramapimod, and the CASP1 inhibitor VX-765. Only VX-765 affected VbP-induced inflammasome activation, whereas all three inhibitors potently blocked the release of IL-1β from poly(I:C) transfected primary keratinocytes (Fig. 4D).

Next, we determined if TAK1 kinase activity is required for NLRP1 inflammasome activation. We found that WT TAK1 but not the kinase inactive mutant TAK1 K63W induced p38 phosphorylation and NLRP1-dependent inflammasome activation in reconstituted 293T cells (Fig. 4E and 4F). Thus, our findings suggest that TAK1 links dsRNA-induced innate immune signaling to the NLRP1 inflammasome in part through its activation of p38.

### Phosphorylation motifs in the NLRP1-DR are required for TAK1-driven NLRP1 inflammasome activation

During RSR-triggered NLRP1 inflammasome activation, ZAKα phosphorylates the NLRP1-DR at two key motifs ^111^PTSTAVL (motif 1) and ^177^PTSTAVL (motif 2) (Fig. 5A)^38^. To determine if TAK1 phosphorylation of the NLRP1-DR at these motifs drives inflammasome activation, we mutated each motif alone or in combination. We found that, consistent with our prior results, NLRP1 responds robustly to poly(I:C) when co-transfected with MDA5 in 293T cells. In contrast, we did not detect inflammasome activation in response to poly(I:C) in 293T cells transfected with variants of NLRP1 in which each motif was mutated individually to alanines (NLRP1^motif1-3A^ and NLRP1^motif2-3A^) or in combination (NLRP1^2×3A^) (Fig. 5B). Similarly, NLRP1^2×3A^ was non-responsive to over-expression of MAVS, TRIF, TAK1, and ZAKα (Fig. 5C). Each NLRP1 phosphomotif mutant was responsive to VbP (Fig. 5B and 5C), highlighting the specificity of N-terminal phosphorylation for MAPK-dependent modes of NLRP1 inflammasome activation, and suggesting that TAK1 also drives inflammasome activation downstream of dsRNA pattern recognition by directly or indirectly phosphorylating the NLRP1-DR.

**Fig. 5.**
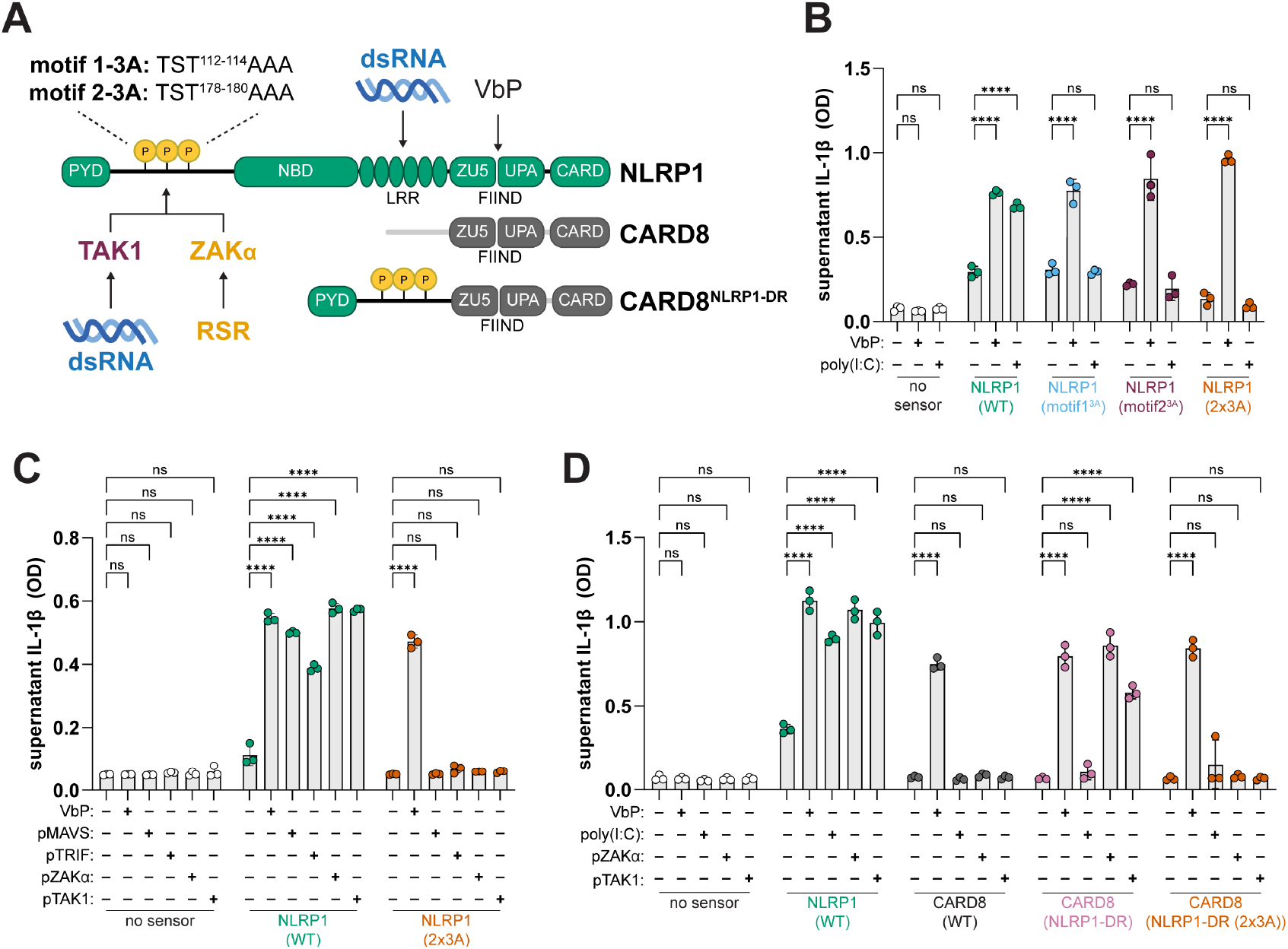
Phosphorylation motifs in the NLRP1 disordered region are required for TAK1-dependent inflammasome activation. (**A**) Schematic depicting phosphorylation motifs in NLRP1’s N-terminal disordered region. (**B-C**) 293T cells were co-transfected with plasmids encoding MDA5 and WT or NLRP1 phophomotif mutants and treated with VbP (10μM) or HMW poly(I:C) (1μg/mL) (B and C) or co-transfected with plasmids encoding MAVS, TRIF, ZAKα, or TAK1 (C). (**D**) 293T cells were co-transfected with plasmids encoding MDA5 and WT NLRP1, WT CARD8, CARD8^NLRP1-DR^, or CARD8^NLRP1-DR2×3A^ and treated with either treated with VbP (10μM) or HMW poly(I:C) (1μg/mL) or co-transfected plasmids encoding ZAKα or TAK1. Supernatant IL-1β levels were measured using an IL-1β bioassay (B-D). Results are representative of two (B and D) or three (C) independent experiments. Data are presented as mean ± SD. p-values were determined by two-way ANOVA with Tukey’s test. *p<0.05, **p<0.01, ***p<0.001, ****p<0.0001.

Based on these observations, we reasoned that phosphorylation of the NLRP1-DR may be sufficient for NLRP1’s response to dsRNA. To test this possibility, we fused the NLRP1 PYD and DR (a.a. 1-254) onto the FIIND–CARD of the related inflammasome-forming sensor CARD8, which is not activated by dsRNA (Fig. 5A)^44^. As expected, WT CARD8 responded to VbP but not ZAKα or TAK1 over-expression. In contrast, we found that the NLRP1 N-terminus endows CARD8 (CARD8^NLRP1-DR^) with the ability to respond to ZAKα and TAK1 in a manner dependent on the presence of phophomotifs 1 and 2. (Fig. 5D). Collectively, our findings suggest that TAK1-mediated phosphorylation of the NLRP1-DR is sufficient to induce inflammasome activation in the absence of exogenous dsRNA (Fig. 5D) and necessary for dsRNA-triggered NLRP1 inflammasome activation (Fig. 5B and 5C). However, poly(I:C) failed to activate the CARD8^NLRP1-DR^ chimeric sensor (Fig. 5D)^38^, consistent with prior findings that dsRNA binding is required for dsRNA-triggered NLRP1 inflammasome activation^44^.

### Physiologic sources of dsRNA induce NLRP1 inflammasome activation

NLRP1 inflammasome activation is induced by arthritogenic alphavirus infection^36,44^. To assess the role of TAK1 in the context of viral infection, we infected N/TERT-2G cells with the Sindbis virus (SINV) strains AR86 and Girdwood (GW) at an MOI of 20 for 48h and measured levels of supernatant IL-1β. Consistent with prior reports demonstrating that viral dsRNA produced during alphavirus infection triggers NLRP1 inflammasome activation^36,44^, we detected elevated levels of IL-1β in the supernatant of SINV-infected WT cells, which was markedly reduced in *NLRP1* KO and *MAP3K7* KO N/TERT-2G cells (Fig. 6A). Moreover, SINV infection of primary keratinocytes also triggered inflammasome activation. In line with our genetic studies in N/TERT-2G cells, SINV-induced inflammasome activation in primary keratinocytes was blocked by TAK1 inhibition (Takinib) to the same extent as treatment with inhibitors of p38 (Doramapimod) and CASP1 (VX-765) (Fig. 6B).

**Fig. 6.**
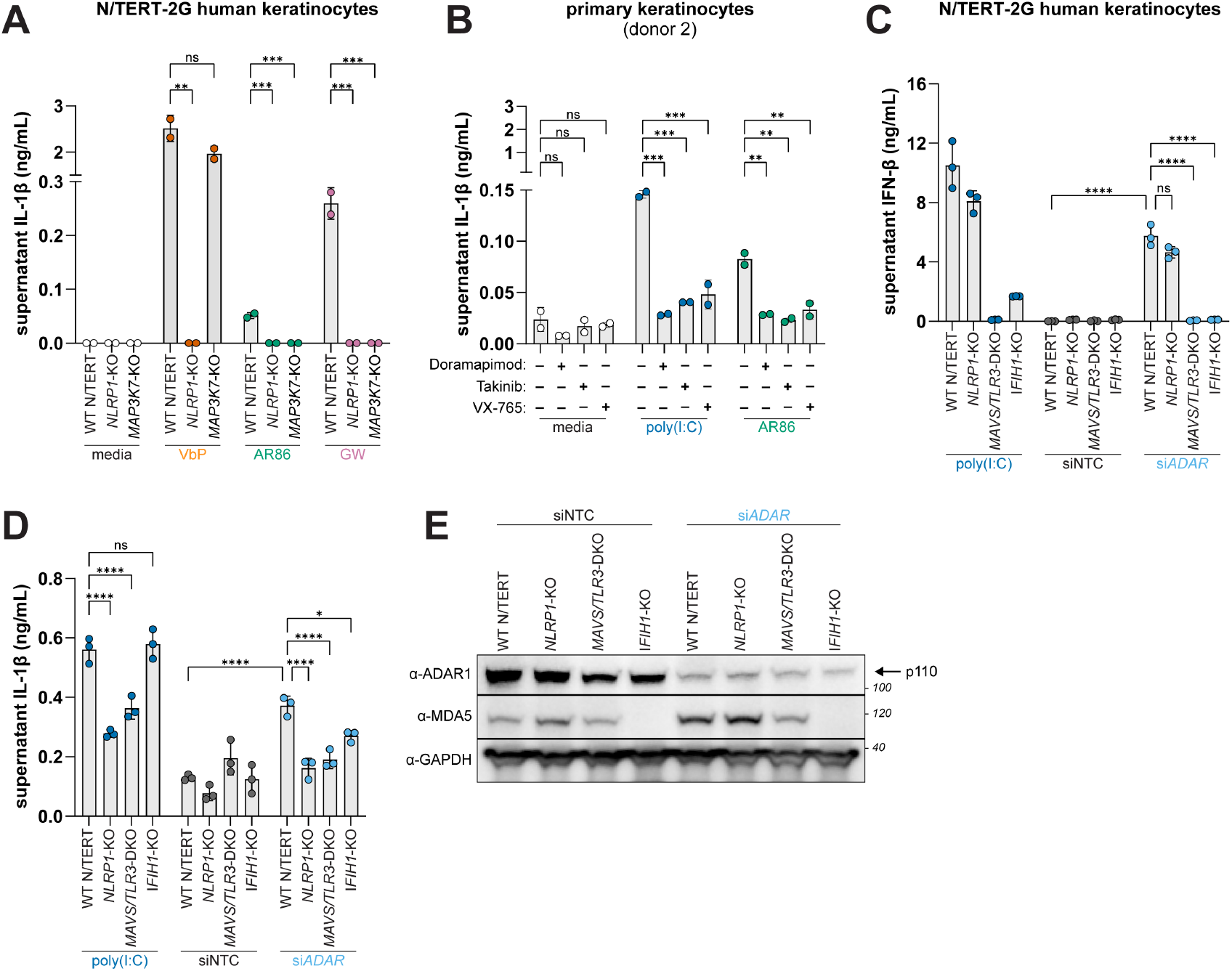
Physiologic sources of dsRNA induce NLRP1 inflammasome activation. (**A**) WT and indicated KO N/TERT-2G cells were treated with either VbP (10μM) or infected with either SINV AR86 or GW (MOI=20) for 48 hours. IL-1β was measured using an antibody-based immunoassay (**B**) Primary keratinocytes were treated with Doramapimod (10μM) or Takinib (10μM) for 2 hours prior to infection with SINV AR86 (MOI=20) for 48 hours. Supernatant IL-1β was measured as in (A). (**C-E**) WT and indicated KO N/TERT-2G cells were treated with siRNAs against *ADAR* or a non-targeting control (NTC) for 24 hours and primed with 10ng/mL TNFα. Supernatants were harvested 72 hours following siRNA treatment and IL-1β (E) and IFN-β (F) levels were measured using an antibody-based immunoassay. Immunoblotting for indicated proteins was carried out as described in (G) from lysates harvested 72 hours post transfection (G). Results are representative of two (A-E) independent experiments. Data are presented as mean ± SD. p-values were determined by one-way ANOVA with Dunnett’s test (A-B) or by two-way ANOVA with Sidak’s test (C-D) using GraphPad PRISM 10. *p<0.05, **p<0.01, ***p<0.001, ****p<0.0001.

Next, we determined if cellular dsRNA can also trigger NLRP1 inflammasome activation in a TAK1-dependent manner. Impairment of Adenosine deaminase acting on RNA 1 (ADAR1) leads to an accumulation of immunostimulatory RNA and is associated with human interferonopathies^56,57^. To model this scenario, we transfected N/TERT-2G cells with small interfering RNAs (siRNAs) targeting *ADAR* or a non-targeting control. Relative to the NTC, *ADAR* knockdown markedly reduced ADAR1 protein levels concomitant with the production of IFN-β in WT and *NLRP1* KO N/TERT-2G cells. Consistent with prior reports^58^, the increase in IFN-β levels by *ADAR1* knockdown was ablated in *IFIH1* (i.e., MDA5) KO and *MAVS*/*TLR3* DKO N/TERT-2G cells (Fig. 6C and 6E), a proxy for the accumulation of unedited dsRNA. Under these conditions, we found that *ADAR1* knockdown also induced IL-1β secretion in WT but not *NLRP1* KO, *IFIH1* KO, or *MAVS*/*TLR3* DKO N/TERT-2G cells (Fig. 6D).

Our findings demonstrate that both viral and cellular dsRNA can trigger NLRP1 inflammasome activation, implicating NLRP1 in the innate immune response to viral infection as well as human autoinflammatory diseases.

## DISCUSSION

dsRNA is an immunostimulatory PAMP produced during viral replication^3,4^. In addition to its well-characterized role in inducing antiviral defenses, including the production of type I IFN, recent studies have demonstrated that dsRNA also activates the NLRP1 inflammasome^36,44,45^. Here we uncover the pathway by which dsRNA triggers MAPK-dependent NLRP1 inflammasome activation. Our findings reveal that NLRP1 inflammasome activation is integrated into dsRNA pattern recognition and signaling by RIG-I/MDA5–MAVS or TLR3– TRIF via the MAP3K TAK1. Below we discuss an emerging model for dsRNA-triggered NLRP1 inflammasome activation and the implications of our findings for NLRP1’s role in the inflammatory response to viral infection as well as human inflammatory diseases associated with dsRNA innate immune signaling.

### Toward a molecular explanation for dsRNA-triggered NLRP1 inflammasome activation

Mechanistically, it remains unresolved how MAPK activity and dsRNA binding cooperate to activate NLRP1. Our findings demonstrate that exogenous dsRNA is not strictly required for NLRP1 inflammasome activation in response to RLR/TLR3 signaling or TAK1 kinase activation. For example, enforced dimerization of the RIG-I CARD and MAVS or TRIF over-expression is sufficient to induce NLRP1 inflammasome activation (Fig. 1G, Fig. 2D). Similarly, TAK1 kinase activity is required for its activation of NLRP1 (Fig. 4D). Moreover, we find that the NLRP1-DR confers responsiveness to ZAKα and TAK1 onto the CARD8 inflammasome in a manner dependent on the presence of MAP3K phosphomotifs in the NLRP1-DR (Fig. 5D). Thus, we posit that phosphorylation rather than dsRNA binding is the key molecular event that triggers NLRP1 inflammasome activation in response to dsRNA.

Our findings do not exclude a role for NLRP1 binding to dsRNA. Indeed, the inability of the CARD8^NLRP1-DR^ chimeric sensor to be activated by poly(I:C) despite its responsiveness to enforced RLR signaling, ZAKα, and TAK1 (Fig. 5D) is consistent with the possibility that dsRNA binding somehow ‘licenses’ NLRP1 inflammasome activation. Additional regulation beyond MAPK activity is appealing as a mechanism to limit NLRP1 inflammasome activation by acute, homeostatic, or otherwise innocuous signaling events. For other liganded inflammasome-forming sensors like NAIP and AIM2, ligand binding induces a conformational change that initiates inflammasome assembly^59,60^. It is possible that NLRP1 binding to dsRNA induces a conformational change that, for example, makes the NLRP1-DR more accessible to TAK1 and p38 phosphorylation; a regulatory step that is seemingly not required for ZAKα-dependent activation^38^. We find this scenario unlikely given that none of the NLRP1 domains that would putatively restrict N-terminal phosphorylation are present in the CARD8^NLRP1-DR^ chimeric sensor (Fig. 5D).

An alternative possibility is that access to the NLRP1-DR or MAP2Ks by ZAKα and TAK1 are distinct in other ways. For example, activated ZAKα has been proposed to disassociate from ribosomes^40^. In contrast, activated TAK1 forms a signalosome initiated by TRAF6 and the TAK1-binding proteins TAB1, TAB2, and TAB3^61^, which may limit its access to NLRP1. We speculate that HMW dsRNA scaffolds NLRP1 with a TAK1-associated signaling complex, bringing the NLRP1-DR into proximity for TAK1 phosphorylation, an otherwise low frequency event. In this model, sustained RLR/TLR3 signaling or TAK1 kinase activity (e.g., in our over-expression studies; Fig 1G and Fig. 4E) bypasses this layer of regulation. A scaffolding model also accounts for the ability of the CARD8^NLRP1-DR^ chimera to respond to ZAKα and TAK1 over-expression but not to poly(I:C) transfection (Fig. 5D), as well as the prior observation that high molecular weight (HMW; >1000bp) dsRNA drives NLRP1 inflammasome activation but low molecular weight dsRNA does not^44^.

We also note that although ZAKα is not activated by dsRNA, the loss of ZAKα nevertheless dampens phospho-p38 levels and dsRNA-triggered NLRP1 inflammasome activation (Fig. 4). Similarly, *MAP3K7* KO N/TERT-2G cells modestly reduce NLRP1 inflammasome activation in response to ribotoxic stress (Fig. 4A). We posit that tonic MAPK activity, as well as regulation by phosphatases (see accompanying manuscripts from Paradis et al. and Chua et al.) also dictates the sensitivity of NLRP1 to phosphorylation-dependent activating stimuli. Ultimately, how phosphorylation drives NLRP1 N-terminal degradation for inflammasome activation remains enigmatic.

### Consequences of NLRP1’s integration into dsRNA innate immune signaling

We and others have previously demonstrated that the NLRP1 inflammasome can directly sense pathogens via the innate immune detection of pathogen-encoded virulence factor activities (e.g., viral protease and E3 ubiquitin ligase enzymatic functions)^18,59,62^. Our findings, along with prior studies characterizing NLRP1 inflammasome activation by ribotoxic stress^35-38^, suggest that NLRP1 is also capable of indirectly sensing pathogens through surveillance of pathogen-associated perturbations that initiate MAP kinase signaling. We speculate that NLRP1’s integration into other stress or innate immune signaling pathways fortify host defenses against pathogen antagonism. For example, NLRP1-dependent pyroptotic death (to eliminate infected cells) or inflammatory signaling (to stimulate cellular immunity) may sustain antiviral defenses triggered by dsRNA even if cellular responses to type I IFN have been inhibited.

The integration of NLRP1 into the innate immune response to dsRNA and our finding that cellular RNA can also drive NLRP1-dependent inflammasome activation also suggests an expanded role for NLRP1 as a driver of viral pathogenesis and autoinflammatory diseases (Fig. 6A-G). Loss of tolerance to nucleic acids and aberrant IFN signaling are established drivers of interferonopathies and other inflammatory diseases associated with spontaneous RLR or TLR3 signaling including psoriasis and lupus, which are also characterized by mucocutaneous pathologies^63,64^. Intriguingly, genetic associations between these diseases and *NLRP1* have been observed but unexplained^41,43,65,66^, and treatment with the IL-1 receptor antagonist Anakinra has been successful in treating a small number of individuals living with SLE^67,68^. Thus, given NLRP1’s capacity to drive skin inflammatory syndromes^41-43,69^, NLRP1 may be a relevant drug target in the setting of interferonopathies and other autoinflammatory syndromes.

## MATERIALS AND METHODS

All reagents, including primers, recombinant DNA, and inhibitors, as well as their sources and other information can be found in **Supplemental Table 1**.

### Cell culture

Human kidney epithelial (293T), embryonic mouse fibroblasts (3T3-J2), HEK-Blue™ IL-1β, HEK-Blue™ IFN-α/β, and BHK cells were cultured in DMEM supplemented with 10% (or 5%, BHKs) FBS, 4.5g/L D-glucose, and 2mM L-Glutamine. N/TERT-2G cells were cultured in Keratinocyte Serum-Free Media supplemented (KSFM) with 2.5μg human epidermal growth factor and 25mg bovine pituitary extract. De-identified primary human keratinocytes were acquired from the University of Pennsylvania Skin Biology and Diseases Resource-based Center (SBDRC) and were cultured in a 1:1 mixture of complete KSFM and Medium 154 supplemented with 200μM CaCl and Human Keratinocyte Growth Supplement (100x) (KC50:50). Primary keratinocyte media was also supplemented with 10,000U/ml Penicillin, 10,000µg/ml Streptomycin, 25µg/ml amphotericin. All cells were maintained at 37°C, and 5% CO2.

### Constructs and cloning

psPAX2, PVSV-G, pRRL-Cas9-puro, pEFBOS-RIG-I and pcDNA human MDA5 were gifts from Daniel Stetson. RIG-I and MDA5 were subcloned into the pQ-PGK-XIP backbone using SbfI and NotI restriction sites. pEFTak-FLAG-MAVS was a gift from Dr. Nandan Gokhale, and MAVS was subcloned into the pQ-PGK-XIP backbone using SbfI and NotI sites. hTLR3-pcDNA3 was a gift from Saumen Sarkar (Addgene plasmid # 32712) and TLR3 was subcloned into pQ-PGK-XIP using SbfI and NotI restriction sites. pcDNA3-TRIF-CFP was a gift from Doug Golenbock (Addgene plasmid # 13644) and was cloned into pQ-PGK-XIP using InFusion HD (Clonetech) pDONR223-MAP3K7 was a gift from William Hahn & David Root (Addgene plasmid # 23693) and MAP3K7 was subcloned into pQ-PGK-XIP using SbfI and NotI restriction sites. Point mutations in MAP3K7 were introduced using overlapping extension PCR. pcDNA4/TO/Strep-HA-ZAK alpha was a gift from Simon Bekker-Jensen (Addgene plasmid # 141193), and ZAKα was subcloned into pQ-PGK-XIP using SbfI and NotI restriction sites. pLKO-puro-CARD8 generated previously^70^. CARD8 was subcloned into pQ-PGK-XIP using SbfI and NotI restriction sites. pRRL-Cas9-neomycin and pRRL-Cas9-blast were synthesized (GenScript). NLRP1 point mutants (motif 1, motif 2, and double motif mutants) and the CARD8^NLRP1-DR^ chimeric sensor (and point mutants) were synthesized (Twist Bioscience) and subcloned into pQ-PGK-XIP using SbfI and NotI restriction sites. Other inflammasome gene constructs were described previously^31,32,70^.

### NLRP1 inflammasome reconstitution in 293T Cells

293T cells were seeded in 24-well format at 1×10^5^ cells/well 24 hours prior to transfection. For NLRP1 inflammasome reconstitution, plasmids encoding for 5ng of NLRP1 or NLRP1 mutants, 2.5ng of ASC, 5ng of CASP1, and 100ng of pro-IL-1β-V5 were co-transfected with indicated associated plasmids and empty pcDNA3 to a total of 500ng of DNA using 1.5μL Mirus Transit-LT1 (Mirus Bio, VWR) per well. 16-18 hours post-transfection, transfected cells were treated with activating stimuli as indicated. Approximately 24 hours post-treatment (40-42 hours post transfection), conditioned supernatants were harvested for HEK-Blue™ assays for cytokine quantification. 293T cells were additionally co-transfected with 50ng of MDA5, 100ng of RIG-I, 25ng of TLR3, 25ng of TRIF, 100ng of MAVS, 30ng of ZAKα, 30ng of TAK1, 10ng of CARD8, or 10ng of CARD8-chimeras where indicated.

### Small interfering RNA knockdown of *ADAR*

N/TERT-2G cells of the indicated genotypes were seeded in 24-well format at 1.5×10^5^ cells/well 24 hours prior to siRNA transfection. ON-TARGETplus non-targeting control pool or *ADAR*-targeting pooled siRNAs were transfected to a final concentration of 25nM using Lipofectamine RNAiMAX according to the manufacturer’s protocol. 24 hours post-transfection, media was replaced with fresh KSFM containing 10ng/mL of TNFα and conditioned supernatants were harvested 48 hours later for cytokine quantification and immunoblotting for knockdown efficiency.

### Lentiviral transduction and generation of knockout cell lines

293T cells were seeded at 8×10^5^ cells/well in 6-well plates the day prior to transfection. Each 6-well was transfected 1μg psPAX2 (packaging vector), 0.5μg pMD2.G (VSV-G envelope vector), and 1μg of either pRRL or pLKO (lentiviral constructs, described below) were mixed with 10μL Mirus Transit-LT1, a 16 hours post transfection, the media of the transfected cells was replaced with KSFM. Conditioned supernatants were harvested at 48- and 72-hours post-transfection and filtered through a 0.45μM filter, supplemented with polybrene (1:1,000), and added to 6-well plates containing N/TERT-2G cells seeded 24 hours prior to transduction at 4×10^5^ cells/well. Cells were transduced by spinfection at 1,200xg for 90 minutes, 32°C. 24 hours post-transduction, virus-containing media was replaced with KSFM supplemented with either blasticidin (50μg/mL) or G418 (100μg/mL) for selection.

Guide RNAs (gRNAs) were designed using either the Synthego Crispr Design Tool or published previously (Table S1). Double-stranded DNA oligos containing the gRNA sequences were cloned into the pRRL-Cas9-blasticidin or pRRL-Cas9-neomycin lentiviral construct using the In-Fusion cloning kit. Lentiviruses were produced and N/TERT-2Gs were transduced as described in the *Lentiviral Transduction*. Cells surviving antibiotic selection were plated for monoclonal selection by limiting dilution. Monoclonal lines were validated as knockouts for the targeted gene by immunoblot and genotyping.

### Inflammasome agonists, inhibitors, and other treatments

Val-boroPro (VbP) and anisomycin (ANS) were resuspended in DMSO. VbP was used at a final concentration of 10μM. ANS was used at a final concentration of 1μM. High molecular weight (HMW) poly(I:C) was resuspended in nuclease-free water and transfected at a final concentration of 1μg/mL using 1.5uL of Lipofectamine LTX per μg of RNA for N/TERT-2G cells or 3μL per ug of RNA of Mirus Transit-LT1 for 293T cells. Recombinant IFN-β (rIFN-β) was resuspended in sterile water and used at a final concentration of 100U/mL. rIFN-β was not removed prior to cytokine quantification. Takinib, Doramapimod, and Ruxolitnib were resuspended in DMSO. Takinib and Doramapimod were both used at a final concentration of 10μM and Ruxolitnib was used at 1μM. Inhibitors were allowed to incubate in treated wells for 1-2 hours prior to treatment with inflammasome activators. A human type I IFN neutralizing antibody mixture (anti-IFN) was used at a final concentration of 1:50 and allowed to incubate for 1-2 hours prior to treatment with inflammasome activators.

### Immunoblotting

Cells were washed with PBS and subsequently harvested with RIPA lysis buffer supplemented with phosphatase (Pierce) and protease (ThermoFisher) inhibitor tablets. Lysates were clarified via centrifugation for 10 minutes at 10,000xg. Samples were denatured by mixing clarified lysate with 6x Laemmli SDS-sample buffer and boiling for 10 minutes at 99°C. Samples were run on 4-12% Bis-Tris gels (ThermoFisher) in 1x MOPS running buffer (ThermoFisher) at 150V for 1 hour. Alternatively, samples were run on 7.5% constant phostag gels (FujiFilm) in Tris-glycine running buffer containing 25mM Tris, 192mM Glycine, and 0.1% SDS at 180V for 90 minutes. Prior to transfer, phostag gels were incubated in transfer buffer containing 10mM EDTA for 20 minutes and washed for 10 minutes in EDTA-free transfer buffer. All gels were transferred to Immobilon-FL PVDF Membrane (Millipore Sigma) at 35V for 2 hours and blocked in tris-buffered saline with 0.05% Tween-20 (TBST) with 3% BSA. Membranes were incubated in 5% BSA TBST containing primary antibody overnight at 4°C. Blots were washed 3x with TBST for 5 minutes and incubated in 3% BSA TBST containing secondary antibody at RT for 1 hour. Washes were repeated as before and immunoblots were developed using SuperSignal West Pico PLUS or Femto Maximum Sensitivity Chemiluminescent Substrates (ThermoFisher) and imaged on (Azure biosystems c600).

### Cytokine quantification

Supernatant IL-1β and type I IFN from 293T cells was quantified using the HEK-Blue IL-1β and IFN-α/β reporter cells (Invivogen), respectively. Briefly, HEK-Blue IL-1β or IFN-α/β cells were plated at 5×10^4^ cells/well in 96-well format in a total volume of 200μL. 50μL of conditioned supernatants from inflammasome-reconstituted 293Ts were added to the reporter cells and allowed to incubate for approximately 24 hours. SEAP levels were determined via colorimetric assay whereby 20μL of conditioned supernatant from HEK-Blue reporter cells was mixed with 180μL of the colorimetric substrate QUANTI-Blue (Invivogen) and allowed to develop for 15-30 minutes at 37°C. Absorbance was measured at OD655 using a CLARIOstar Plus plate reader and absolute levels of the respective cytokines were calculated relative to the standard curve.

To quantify secreted IL-1β and IFN-β from N/TERT-2G cells, supernatants were harvested from treated N/TERT-2Gs and frozen at −20°C until processing. Samples were diluted 1:1 in KSFM and human IL-1β (R&D Systems) and human IFN-β (R&D Systems) were performed according to the manufacturer’s protocol. Absorbance was measured at 450nm and 570nm using a CLARIOstar Plus plate reader (BMG Labtech). To quantify IL-1β and type I IFN from alphavirus infected N/TERT-2G cells or where indicated, samples were processed in 384-well format using Lumit IL-1β and IFN-β human immunoassay (Promega) according to the manufacturers protocol, which has a larger dynamic range and increased specificity for IL-1β p17 than ELISAs.

### RNA isolation and RT-qPCR

N/TERT-2G cells were rinsed with PBS then lysed using the Power SYBR Green Cells-to-CT Kit (ThermoFisher) lysis buffer according to the manufacturer’s protocol. Reverse transcription and qPCR were performed on a CFX384 Touch Real-Time PCR detection system (Bio-Rad) using the PowerUp SYBR Green Master Mix for qPCR (ThermoFisher) following the manufacturer’s protocol. Probes for *MX1, ISG15, IFIT1*, and *18S* were selected from published works where cited and synthesized using Integrated DNA Technologies (IDT).

### Viral infection and quantification

The titer of viral stocks of the Sindbis virus strains AR86 and Girdwood (GW) were determined by plaque assay. Briefly, 5×10^4^ BHK cells were seeded into 24-well plate approximately 24 hours prior to infection. Supernatants from low MOI (0.1) infections were harvested 48 hours post infection and serially diluted 10-fold in serum-free DMEM. Approximately 24 hours post-seeding, media was removed from BHK-containing wells, and 200μL of serially diluted supernatant was overlayed and incubated for approximately 1 hour. Following infection, each well received 500uL of overlay containing a 1:1 mixture of 2% methylcellulose (4,000 cP) and complete growth media (final methylcellulose concentration of 1%). Infections were allowed to progress under overlay for 5 days. On day 5, cells were fixed using 10% formaldehyde in PBS for 2 hours. The overlay and fixative was aspirated, and wells were stained using 0.1% crystal violet and 20% methanol for approximately 30 minutes. Stained wells were washed gently using de-ionized water and were counted for PFU.

AR86 and GW were resuspended in 50µL KSFM or KC50:50 and added to 1.5 × 10^5^ N/TERT-2G cells or primary keratinocytes in 24-well plates respectively. All cells were infected at an MOI of 20 and supernatants were harvested after 48 hours.

## Statistical Analysis

Statistical analyses were performed using GraphPad Prism 10.

## Supplementary Materials

Fig. S1-S3

Table. S1

## Acknowledgments

We thank Matt Daugherty, Adam Geballe, and Tristan Jordan for critical feedback on the manuscript, Veit Hornung, Etienne Meunier, and Franklin Zhong for helpful discussions and the sharing of unpublished data, Andrew Oberst, Ram Savan, Daniel Stetson, Cory Simpson, and members of their labs for sharing reagents and protocols, and all members of the Mitchell lab for input on the project. This work was supported by the University of Pennsylvania Skin Biology and Diseases Resource-based Center (SBDRC). PSM is a Freeman Hrabowski Scholar of the Howard Hughes Medical Institute.

## Funding

National Institutes of Health NRSA T32GM136534 (HCT, MAY, MRC)

National Institutes of Health grant DP2AI154432 (PSM)

National Institutes of Health grant P30AR069589 (PSM via the UPENN SBDRC)

Howard Hughes Medical Institute (PSM)

## Author contributions

Conceptualization: MRC, PSM

Methodology: MRC, PSM, JLH

Investigation: MRC, AH, JLH, HCT, MAY, AC

Visualization: MRC, PSM

Funding acquisition: MRC, PSM

Project administration: PSM

Supervision: PSM

Writing – original draft: MRC, PSM

Writing – review & editing: MRC, PSM, AH, JLH, HCT, MAY, AC

## Data and materials availability

All data are available in the main text or the supplementary materials.

