## Supplementary Materials for "TAK1 integrates the NLRP1 inflammasome into the innate immune response to double-stranded RNA"

Fig. S1-S3

Table. S1

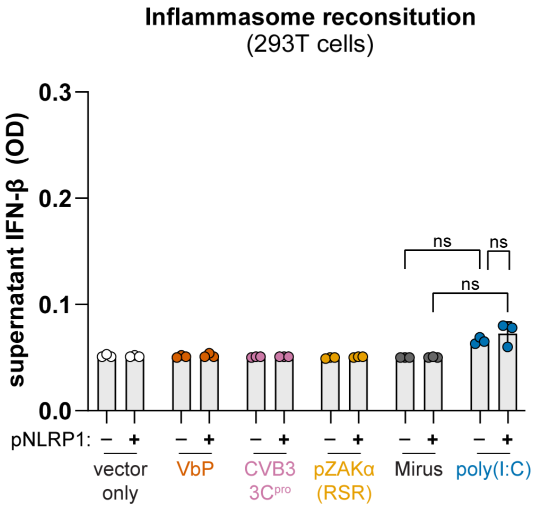


**Supplementary Figure 1. dsRNA does not induce type I IFN production in 293T cells.** Data correspond to Fig. 1A and 1B. Reconstitution of the NLRP1 inflammasome by co-transfecting 293T cells with plasmids encoding NLRP1, ASC, CASP1, and pro-IL-1β. Cells were treated with VbP (10μM), mock transfected, or co-transfected with HMW poly(I:C) (1μg/mL) or plasmids encoding ZAKα or CVB3 3C protease. Bioactive IFN-β levels were quantified using a HEK-Blue IFN reporter assay. Data are presented as mean ± SD. p-values were determined by two-way ANOVA. *p<0.05, **p<0.01, ***p<0.001, ****p<0.0001.

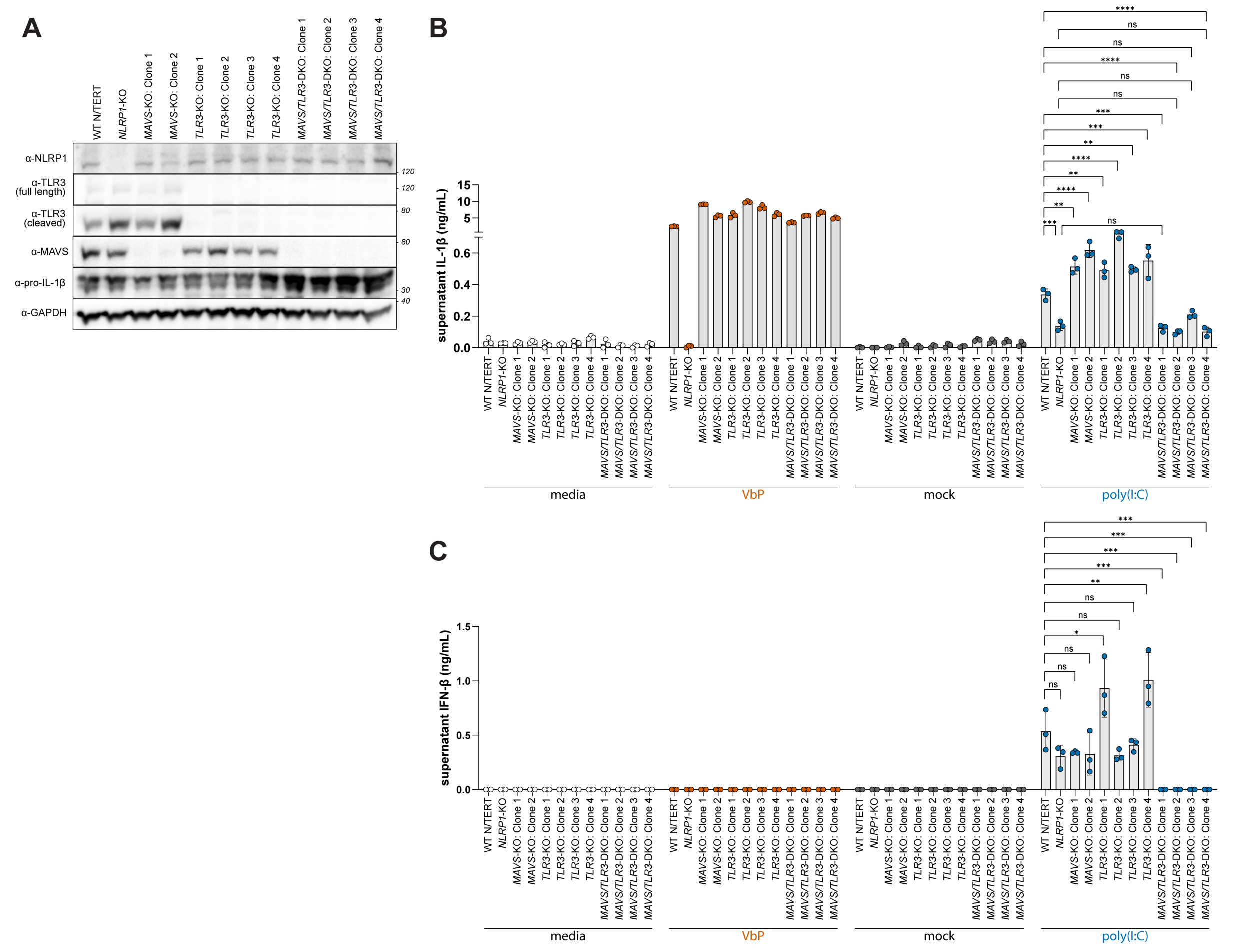


**Supplementary Figure 2. Validation of N/TERT-2G knockout cell lines associated with Fig 2.**
(**A**) Immunoblotting for indicated proteins using lysates generated from N/TERT-2G monoclonal KO cell lines characterized in Fig. 2. (**B-C**) Expanded data associated with Fig. 2A-B. Data are presented as mean ± SD. p-values were determined by one-way ANOVA with Dunnett's test (A-B). *p<0.05, **p<0.01, ***p<0.001, ****p<0.0001.

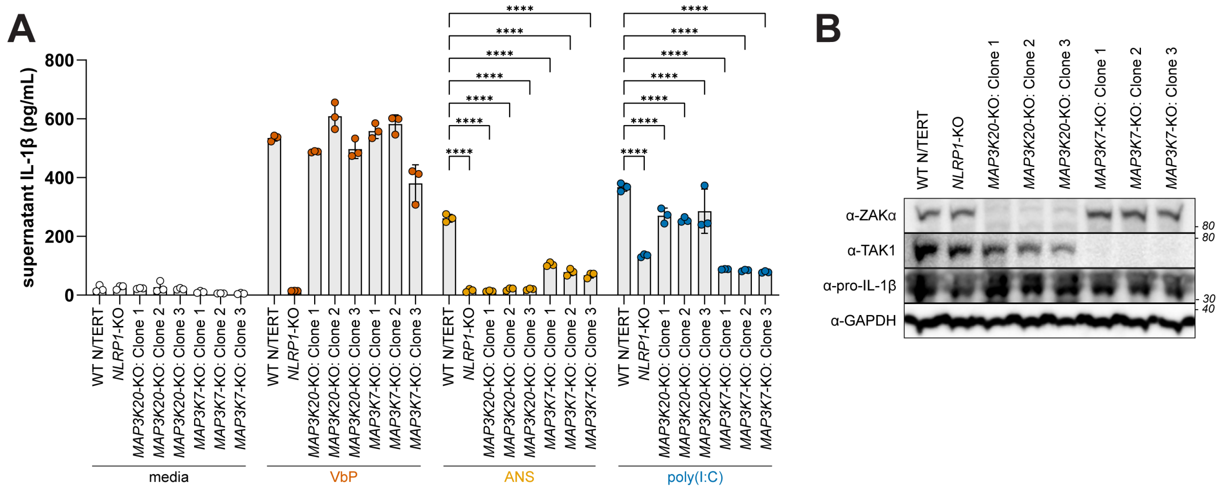


**Supplementary Figure 3. Validation of N/TERT-2G knockout cell lines associated with Fig 4.**
**(A)** Expanded data is associated with Fig. 4A. (**B**) Immunoblotting for indicated proteins using lysates generated from N/TERT-2G monoclonal knockout (KO) cell lines characterized in Fig. 4. Data are presented as mean ± SD. p-values were determined by two-way ANOVA with Sidak's test. *p<0.05, **p<0.01, ***p<0.001, ****p<0.0001

**Supplemental Table S1. Reagents used in this study.** We provide the source, manufacturer, and other information for key reagents, including antibodies, cell lines, and inhibitors, used in this study.

| REAGENT or RESOURCE | SOURCE | IDENTIFIER |
|---|---|---|
| Antibodies | | |
| MAVS Antibody (1:500) | Cell Signaling Technologies | Cat# 3993S |
| Toll-like Receptor 3 (D10F10) (1:500) | Cell Signaling Technologies | Cat# 6961S |
| Phospho-p38 MAPK (Thr180/Tyr182) (D3F9) Rabbit Monoclonal Antibody (1:500) | Cell Signaling Technologies | Cat# 4511S |
| Human IL-1 beta /IL-1F2 Antibody (1:250) | R&D Systems | Cat# AF-201-NA |
| TRIF Antibody (1:1000) | Cell Signaling Technologies | Cat# 4596S |
| TAK1 (D94D7) Rabbit mAb (1:500) | Cell Signaling Technologies | Cat# 5206S |
| V5-Tag (D3H8Q) (1:1000) | Cell Signaling Technologies | Cat# 13202S |
| ZAK antibody (1:500) | Fortis Life Sciences | Cat# A301-993A |
| p38 MAPK Antibody | Cell Signaling Technology | Cat# 9212S |
| GAPDH (14C10) Rabbit mAb (1:1000) | Cell Signaling Technologies | Cat# 2118S |
| Human NLRP1/NALP1 Antibody (1:250) | R&D Systems | Cat# AF6788 |
| ADAR Monoclonal Antibody (GT1066) (1:500) | ThermoFisher | Cat# MA5-17285 |
| MAVS Antibody (1:500) | Cell Signaling Technologies | Cat# 3993S |
| Rig-I (D14G6) Rabbit mAb (1:500) | Cell Signaling Technologies | Cat# 3743S |
| MDA-5 (D74E4) Rabbit mAb (1:500) | Cell Signaling Technologies | Cat# 5321S |
| IRF-3 (D83B9) Rabbit mAb (1:500) | Cell Signaling Technologies | Cat# 4302S |
| Monoclonal ANTI-FLAG M2 antibody (1:1000) | Millipore Sigma | Cat# F3165-1MG |
| Goat anti-Mouse IgG (H+L) Poly-HRP Secondary Antibody, HRP (1:10000) | VWR | Cat# 10147-306 |
| Goat anti-Rabbit IgG (H+L) Poly-HRP Secondary Antibody, HRP (1:10000) | VWR | Cat# 10147-304 |
| Donkey anti-Sheep IgG (H+L) Secondary Antibody, HRP | Invitrogen | Cat# A16041 |
| Bacterial and virus strains | | |
| SINV AR86 | Provided by Dr. Jennifer Hyde | N/A |
| SINV Girdwood | Provided by Dr. Jennifer Hyde | N/A |
| Biological samples | | |
| Primary Keratinocyte: Donor 5230 | University of Pennsylvania Skin Biology and Diseases Resource-based Center (UPenn SBDRC) | N/A |
| Primary Keratinocyte: Donor 5231 | UPenn SBDRC | N/A |
| Primary Keratinocyte: Donor 5240 | UPenn SBDRC | N/A |
| Chemicals, peptides, and recombinant proteins | | |
| Human IL-1 beta Recombinant Protein | Invitrogen | Cat# A42509 |
| Interferon-β Protein, Recombinant human | Millipore Sigma | Cat# IF014 |
| Recombinant human TNF-α | Invivogen | Cat# rcyc-htnfa |

| | | |
|---|---|---|
| Human Type I IFN Neutralizing Antibody Mixture | PBL Assay Science | Cat# 39000 |
| Takinib | SelleckChem | Cat# S8663 |
| Doramapimod | SelleckChem | Cat# S1574 |
| Ruxolitinib | MedChemExpress | Cat# HY-50856 |
| VX-765 | InvivoGen | nh-vx765i-1 |
| Polybrene Infection / Transfection Reagent | Millipore Sigma | Cat# TR-1003-G |
| Val-boroPro | Millipore Sigma | Cat# 5314650001 |
| TransIT-LT1 Transfection Reagent, Mirus Bio | VWR | Cat# 10767-122 |
| Lipofectamine RNAiMAX Transfection Reagent | ThermoFisher | Cat# 13778075 |
| Lipofectamine LTX | ThermoFisher | Cat# 15338100 |
| Anisomycin | ThermoFisher | Cat# 50-187-3712 |
| Poly(I:C) HMW | InvivoGen | Cat# tlrl-pic |
| ON-TARGETplus Human ADAR siRNA (SMARTPool) | HorizonDiscovery | Cat# L-008630-00-0005 |
| N-TARGETplus Non-targeting Control Pool | HorizonDiscovery | Cat# D-001810-10-05 |
| Critical commercial assays | | |
| Human IL-1 beta/IL-1F2 DuoSet ELISA | R&D Systems | Cat# DY201 |
| Human IFN-beta DuoSet ELISA | R&D Systems | Cat# DY814-05 |
| DuoSet ELISA Ancillary Reagent Kit 2 | R&D Systems | Cat# DY008B |
| Power SYBR™ Green Cells-to-CT™ Kit | ThermoFisher | Cat# 4402953 |
| PowerUp™ SYBR™ Green Master Mix for qPCR | ThermoFisher | Cat# A25742 |
| In-Fusion® Snap Assembly Master Mix | Takara Biosciences | Cat# 638947 |
| Lumit® IFN-β (Human) Immunoassay | Promega | Cat# W1810 |
| Lumit® IL-1β Human | Promega | Cat# W6011 |
| Cell lines | | |
| HEK293T | N/A | RRID:CVCL_0063 |
| N/TERT-2G | Provided by Dr. Cory Simpson; PMID: 10648628 | RRID:CVCL_UZ62 |
| BHK | Provided by Dr. Jennifer Hyde | RRID:CVCL_1915 |
| HEK-Blue™ IL-1β | InvivoGen | Cat# hkb-il1bv2; RRID:CVCL_A8CL |
| HEK-Blue™ IFN-α/β | InvivoGen | Cat# hkb-ifnabv2; RRID:CVCL_KT26 |
| Oligonucleotides | | |
| qPCR-MX1-F: TTCAGCACCTGATGGCCTATC | Holzinger et al. 2007; PMID: 17494065 | N/A |
| qPCR-MX1-R: TGGATGATCAAAGGGATGTGG | Holzinger et al. 2007: PMID: 17494065 | N/A |
| qPCR-IFIT1-F: TGGTGACCTGGGGCAACTTT | Otter and Bracci et al. 2024; PMID: 38568967 | N/A |
| qPCR-IFIT1-R: AGGCCTTGGCCCGTTCATAA | Otter and Bracci et al. 2024; PMID: 38568967 | N/A |
| qPCR-ISG15-F: CATCTTTGCCAGTACAGGAGC | Otter and Bracci et al. 2024; PMID: 38568967 | N/A |
| qPCR-ISG15-R: GGGACACCTGGAATTCGTTG | Otter and Bracci et al. 2024; PMID: 38568967 | N/A |
| qPCR-18S-F: TTCGATGGTAGTCGCTGTGC | Otter and Bracci et al. 2024; PMID: 38568967 | N/A |
| qPCR-18S-R: CTGCTGCCTTCCTTGAATGTGGTA | Otter and Bracci et al. 2024; PMID: 38568967 | N/A |
| gRNA-*MAVS*: GAGGTGGCCCGCAGTCGATCC | Synthego | N/A |
| gRNA-*IRF3*: GGCCACGGATCCTGCCCTGGC | Zhu et al. 2024; PMID: PMID: 39362857 | N/A |
| gRNA-*MAP3K7*: GAGTTGTTTGCAAAGCTAAG | Synthego | N/A |
| gRNA-*MAP3K20#1*: CATTAACAGTAACAGAAGTG | Synthego | N/A |
| gRNA-*MAP3K20#2*: GATATGGATCACATTATGACC | Synthego | N/A |
| gRNA-*TLR3#1*: CCAGGTCAAGTACTTCTAGG | Synthego | N/A |
| gRNA-*TLR3#2*: GTCAACGACTGATGCTCCGAA | Synthego | N/A |
| gRNA-*IFIH1*: GTGAGAAAGAAAGATGTCGAA | Synthego | N/A |
| gRNA-*NLRP1*: TGTAGGGGAATGAGGGAGAG | Synthego | N/A |

| | | |
|---|---|---|
| Recombinant DNA | | |
| psPAX2 | Provided by Dr. Dan Stetson; PMID: 25064072 | N/A |
| pVSV-G | Provided by Dr. Dan Stetson; PMID: 25064072 | N/A |
| pRRL-Cas9-Puro | Provided by Dr. Dan Stetson; PMID: 25064072 | N/A |
| pRRL-Cas9-Blast | This paper | N/A |
| pRRL-Cas9-Neomycin | This paper | N/A |
| pEFBOS-RIGI | Provided by Dr. Dan Stetson | N/A |
| pQ-PGK-XIP-RIG-I | This paper | N/A |
| pCDNA3 human MDA5 | Provided by Dr. Dan Stetson | N/A |
| pQ-PGK-XIP-MDA5 | This paper | N/A |
| pEFTak FLAG-MAVS | Provided by Dr. Nandan Gokhale; PMID: 39700280 | N/A |
| pQ-PGK-XIP-MAVS | This paper | N/A |
| hTLR3-pcDNA3 | Addgene; PMID: 20382888 | Cat# 32712 |
| pQ-PGK-XIP-TLR3 | This paper | N/A |
| pcDNA3-TRIF | Addgene, Doug Golenbock, Unpublished | Cat# 13644 |
| pQ-PGK-XIP-TRIF | This paper | N/A |
| pDONR223-MAP3K7 | Addgene; PMID: 21107320 | Cat# 23693 |
| pQ-PGK-XIP-MAP3K7 (TAK1) | This paper | N/A |
| pQ-PGK-XIP- MAP3K7 (TAK1)-K63W | This paper | N/A |
| pcDNA4/TO/Strep-HA-ZAK alpha | Addgene; PMID: 32289254 | Cat# 141193 |
| pQ-PGK-XIP-ZAKα | This paper | N/A |
| pQ-PGK-XIP-NLRP1 | This paper | N/A |
| pQ-PGK-XIP-NLRP1-3A (T112A/S113A/T114A) | This paper | N/A |
| pQ-PGK-XIP-NLRP1-3A (T178A/S179A/T180A) | This paper | N/A |
| pQ-PGK-XIP-NLRP1-2x3A (T112A/S113A/T114A/T178A/S179A/T180A) | This paper | N/A |
| pQ-PGK-XIP-CARD8 | This paper | N/A |
| pQ-PGK-XIP-CARD8$^{NLRP1\text{-}DR}$ | This paper | N/A |
| pQ-PGK-XIP-CARD8$^{NLRP1\text{-}DR\text{-}2x\text{-}3A}$ | This paper | N/A |
| pRRL 3xFV-N-RIG-I$^{CARD}$ | Provided by Dr. Lan Chu; PMID: 39700280 | N/A |
| Inflammasome reconstitution plasmids: human ASC, human CASP1, and human IL-1B-V5 | PMID: 27926929 | N/A |
| Software and algorithms | | |
| GraphPad Prism 10 | GraphPad Software | RRID:SCR_002798 |
| Geneious | Geneious | RRID:SCR_010519 |
| Synthego Crispr Design Tool | Synthego | RRID:SCR_026304 |
| Integrated DNA Technologies (IDT) | Integrated DNA Technologies (IDT) | RRID:SCR_025813 |
| Immunoblotting | | |
| MagicMark™ XP Western Protein Standard | ThermoFisher | Cat# LC5602 |
| iBright™ Prestained Protein Ladder | Thermofisher | LC5615 |
| SuperSignal™ West Pico PLUS Chemiluminescent Substrate | ThermoFisher | Cat# 34577 |
| SuperSignal™ West Pico Femto Chemiluminescent Substrate | ThermoFisher | Cat# 34085 |
| SuperSep Phos-tag (50μmol/l), 7.5%, 17well, 100×100×6.6 mm | FujiFilm | Cat# 192-18001 |
| Bolt™ 4 to 12%, Bis-Tris, 1.0 mm, Mini Protein Gels, 15-well | ThermoFisher | Cat# NW04125 |
| NuPAGE™ MOPS SDS Running Buffer (20X) | ThermoFisher | Cat# NP0001 |
| Immobilon®-FL PVDF Membrane | Millipore Sigma | Cat# IPFL00010 |
| Pierce™ Protease Inhibitor Mini Tablets, EDTA-free | Thermofisher | Cat# A32955 |
| Pierce™ Phosphatase Inhibitor Mini Tablets | Thermofisher | Cat# A32957 |

| | | |
|---|---|---|
| RIPA Lysis Buffer, 10X | Millipore Sigma | Cat# 20-188 |
| Cell Culture | | |
| Keratinocyte SFM (1X) | ThermoFisher | Cat# 17005042 |
| Human Keratinocyte Growth Supplement (HKGS) | ThermoFisher | Cat# S0015 |
| Medium 154CF, Kit | ThermoFisher | Cat# M154CF500 |
| GlutaMAX Supplement | ThermoFisher | Cat# 35050061 |
| FBS select grade CR origin 500mL HI batch 024A23 | VWR | Cat# 76419-588 |
| Penicillin-Streptomycin (10,000 U/mL) | ThermoFisher | Cat# 15140122 |
| DMEM | Life Technologies | Cat# 11960069 |
| PBS 7.4 1X | ThermoFisher | Cat# 10010049 |
| Trypsin-EDTA (0.5%), no phenol red | ThermoFisher | Cat# 15400054 |
| Trypsin-EDTA (0.05%), phenol red | ThermoFisher | Cat# 25300120 |
| Amphotericin B | ThermoFisher | Cat# 15290026 |
